# Microtubules ensure transport of vegetative nuclei and sperm cells by fine-tuning their home positions

**DOI:** 10.1101/2024.01.31.578224

**Authors:** Kazuki Motomura, Haruna Tsuchi, Marin Komojiri, Ayumi Matsumoto, Naoya Sugi, Daichi Susaki, Atsushi Takeda, Tetsu Kinoshita, Daisuke Maruyama

## Abstract

The pollen tube plays a pivotal role in double fertilization by delivering sperm cells (SCs) to the ovule. In *Arabidopsis thaliana*, a pair of SCs tightly connects with the vegetative nucleus (VN) to form the male germ unit (MGU), which is located in the apical region during pollen-tube growth, keeping the VN ahead of the SCs. MGU transport relies on independent motility of VN and SC pairs. However, the complexity of this dual motive force has hindered our understanding of MGU behavior, including its positioning and nuclear order. We used *Arabidopsis* mutants or transgenic plants that produced semi-motile MGUs with aberrant VNs or SCs to analyze the positioning of VN or SCs after stochastically disconnecting the MGU. In pollen tubes with an immotile SC pair, the VN was ∼70 μm away from the tip, whereas in pollen tubes with an immotile VN, the SC pair was ∼100 μm away from the tip, implying that the VN and SCs have independent home positions. The position of MGU moved forward owing to the loss of the microtubule-destabilizing kinesin *KINESIN-13A*. Conversely, microtubule depolymerization by oryzalin treatment or introducing mutations in *TUBULIN BETA 4* (*TUB4*) deregulated the position of the MGU and shifted its position backward. In addition, *tub4* plants exhibited reduced fertility. These data indicate a significant role of microtubules in stable MGU positioning to ensure reproductive success.

## Introduction

The pollen tube is a tip-growing tissue that delivers the male genome to an ovule during sexual reproduction in flowering plants. In *Arabidopsis thaliana* and other plants that produce tricellular pollen, pollen tube vegetative cells contain two sperm cells (SCs), one of which has a tail-like protrusion tightly associates with the vegetative nucleus (VN) to form a reproductive cell complex known as the male germ unit (MGU). Although MGU vigorously moves back and forth, it gradually moves forward in the net movement accompanying the tip growth of the pollen tube.

In the past decade, genetic evidence of independent motive forces in VN and SC pairs has been shown in *A*. *thaliana*. Loss-of-function of two WPP domain-interacting tail-anchored protein 1 (WIT1) and WIT2 causes passive migration of VN dragged by the SC pair (Zhou and Meier, 2014). WIT1 and WIT2 are localized on the outer nuclear envelope and redundantly serve as components of the linker of the nucleoskeleton and cytoskeleton (LINC) complex (Tatout et al., 2014; Groves et al., 2020). The LINC complex regulates nuclear migration in plant cells by transmitting the mechanical force generated by motor proteins. Loss of other LINC components resulted in similar VN transport defects, indicating a crucial role of the LINC complex in VN motility (Zhou et al., 2015). Although the key molecules involved in SC motility have not yet been identified, SC motility is susceptible to alterations in the extracellular environment of the SC pairs. When the overaccumulation of callose (β-1,3-glucan) was ectopically induced by the *pHTR10:cals3m* transgene (designated *SC-cal*), which expresses callose synthase (*cals3m*) from the SC-specific *HTR10* promoter, SC pairs were observed at the basal region of the pollen tube (Motomura et al., 2021). In these aberrant pollen tubes, MGU connections tend to be lost, and VNs are often detected in the apical region, suggesting loss of SC-specific motility in pollen tubes from *SC-cal* transformants (Motomura et al., 2021).

Active transport of organelles requires cables and motors (Perico and Sparkes, 2018). Pollen tubes contain long actin cables traversing the shank region that are likely to orient their plus end toward the tip, because the actin polymerization factor formin is localized at the tip of the pollen tube (Cheung and Wu, 2004; Cheung et al., 2010). In pollen tubes, various organelles move along actin cables using myosin XI family proteins (Madison et al., 2015). Myosin XI-I/KAKU1 is the only myosin XI associated with the nuclear envelope in *Arabidopsis* (Avisar et al., 2009). Myosin XI-I/KAKU1 controls the spindle shape of the nucleus and LINC-dependent nuclear movement in vegetative tissues (Tamura et al., 2013). However, unlike *wit1 wit2* double mutant plants, male fertility was not reduced in *kaku1* plants (Zhou and Meier, 2014). Overall, myosin function in apical MGU transport is unclear.

The role of microtubules in nuclear transport is unclear, probably because of relatively moderate abnormalities in MGU transport after treatment with microtubule-disrupting drugs (Cai et al., 1997). Immunostaining analysis of tubulins and the GFP-tagged plus-end binding protein AtEB1 showed that pollen tubes contain longitudinally aligned microtubules in the shank region and shorter cables in the subapical region (Cheung et al., 2008; Cai and Cresti, 2010). Kinesins with the calponin homology domain (KCH), which belong to the kinesin 14 subfamily expressed in pollen tubes, are candidates for motor proteins that drive SCs, in a “tug-of-war” model (Cai et al., 1997; Schattner et al. 2021; Cai, 2022). *In vitro* experiments have demonstrated that rice *Os*KCH1 interacts with both microtubules and actin fibers to drive fast (average 69.17 ± 14.77 nm s^−1^) and slow (average 24.2 ± 13.42 nm s^−1^) sliding movements (Schattner et al., 2021; Walter et al., 2015). Because fast and slow movements appeared simultaneously in actin fiber fragments railing the same microtubule, the two velocities were assumed to reflect different motive forces in the parallel and antiparallel sliding of the two fibers. In the tug-of-war model, KCH1 generates different velocities of back-and-forth movements by associating on one side with randomly oriented microtubules surrounding the SCs, and on the other side with uniformly directed long actin cables (Schattner et al., 2021). Although this model provides a simple explanation based on cytological molecular data, there is lack of genetic evidence.

Analysis of the apical MGU transport mechanism was hindered by the behavioral interaction between VN and the SC pair, based on their independent motive force and physical connection through a tail-like protrusion. Previously, we genetically dissected the apical transport of the VNs and SCs (Motomura et al., 2021). *SC-cal* transgenic (immotile SCs) and *wit1 wit2* (immotile VN) plants showed stochastic disconnection of MGU, producing mobile “orphan” VNs and SC pairs, respectively (Motomura et al., 2021). The MGU disconnections opened the door to separately monitor behaviors of VNs and SC pairs, and these plants thus can be key genetic tools to analyze molecular mechanism of the apical MGU transport. Here, we demonstrate that the orphan SC pair in *wit1 wit2* plants was at positions distant from pollen-tube tips (∼100 μm) than orphan VNs in *SC-cal* (∼70 μm), suggesting that VNs and SCs intrinsically determine their own “home positions.” Different home positions in the VN and SC pairs could explain the stable MGU order during pollen-tube growth, because the home positions of each orphan MGU component almost corresponded to those in wild-type MGUs. We found that microtubules and KINESIN-13A (KIN13A) were involved in MGU positioning. These data elucidate the basis of a stable MGU positioning system required for double fertilization.

## Results

### VN and the SC pair have different home positions in the pollen tube

To examine the behavior of a VN separated from its SC pair, we analyzed *in vitro*-germinated pollen tubes from transgenic plants carrying the *SC-cal* transgene and *pRPS5A:HISTONE H2B-tdTomato* (*RHT*) nuclear marker. In this genetic background, approximately 80% of tdTomato-labeled MGUs display disconnection of the VN–SC linkage (Motomura et al., 2021). Apically transported orphan VNs were detected 61.4 ± 29.6 μm away from pollen-tube tip (mean ± SD, Fig. 1, A and B). We analyzed SC pairs separated from the immotile VN *in vitro*-germinated *wit1 wit2* pollen tubes carrying *RHT*. Similar to VN in *SC-cal* plants, orphan SC pairs were detected in the apical region (Fig. 1, C and D, Zhou and Meier, 2014; Motomura et al., 2021). However, orphan SC pairs were located 95.9 ± 42.6 μm away from the tip of *wit1 wit2* pollen tubes, which was much basal than those of orphan VN in *SC-cal* plants (Fig. 1, B and D). As previously reported, we did not observe significant differences in pollen-tube length between wild-type *RHT*, *SC-cal RHT*, and *wit1 wit2 RHT* plants, suggesting that differences in the positioning of VN and SC pair are not associated with the growth activity of pollen tubes (Supplemental Fig. S1) (Motomura et al., 2021). Thus, the VN and SC pair intrinsically determine their own “home positions.” Interestingly, the home positions of the VN and SC pairs were considerably close to those of intact MGUs in wild-type *RHT* plants, implying that the presence of a tail-like connection has a limited effect, at least on the control of the home position (Fig. 1, A to D).

**Figure 1.**
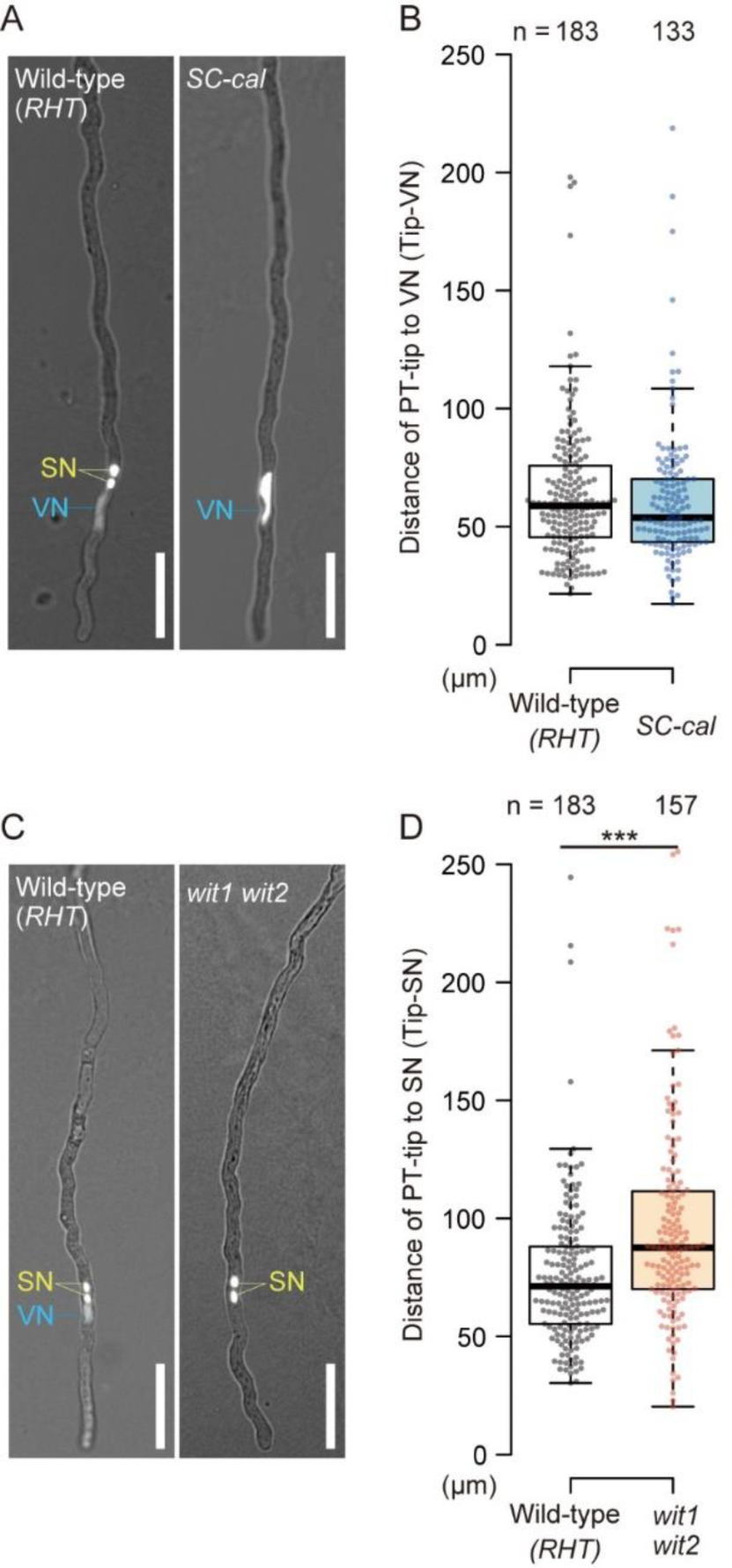
The vegetative nucleus and sperm cell pair have different home positions in the pollen tube. **A)** Representative images of *in vitro*-germinated pollen tubes from wild-type and *SC-cal* transgenic plants carrying the *pRPS5A:HISTONE H2B-tdTomato* (*RHT*) nuclear marker. **B)** Distance from pollen tube-tip to the vegetative nucleus (Tip-VN) analyzed in (**A**). **C)** Representative images of *in vitro*-germinated pollen tubes from wild-type and *wit1 wit2* plants carrying *RHT*. **D)** Distance from pollen tube-tip to sperm nuclei (Tip-SN) analyzed in (**C**). In (**A**) and (**C**), red fluorescent signals were observed with transmitting light 3 h after pollination. Box- and-whisker plots show median (center line), upper and lower quartiles (box), maximum and minimum (whiskers), and points (solid circles). Asterisks, significance by Welch’s *t* test (\*\*\**P* = 8.4 × 10^−6^). Scale bars, 50 μm.

### Microtubule-disrupting drugs disturb positional control of VN and the SC pair

To examine the mechanisms controlling the home positions of the VN and SC pairs, we analyzed the requirement of microtubules under semi-*in vivo* pollen-tube growth conditions. Pollinated pistils were incubated on media with or without oryzalin (a microtubule disrupter) and pollen tubes emerging from the cut style were observed by microscopy. In *SC-cal RHT*, orphan VN was detected approximately 100 μm from the tip in the control medium; however, orphan VN was broadly distributed in pollen tubes treated with 1 μM oryzalin (Fig. 2, A and B). The positional control of orphan SC was compromised in *wit1 wit2 RHT* (Fig. 2, C and D). These data suggest microtubules play an important role in the positional control of the VN and SC pairs. To observe how disturbing the home positions of VN and the SC pair affects MGU behavior, we analyzed wild-type *RHT* pollen tubes under semi-*in vivo* growth conditions. Consistent with the observations for the orphan VN and SC pairs, the positions of VN and SN pairs in wild-type shifted backward and were broadly distributed in the apical region of pollen tubes on oryzalin-containing medium (Fig. 3, A to C). In approximately 40% of the pollen tubes, the order of the MGU was inverted after oryzalin treatment, compared with the control, in which VN was ahead in 90% of the pollen tubes (Fig. 3D). These observations confirmed the significance of microtubules in stabilizing the MGU structure, including the VN-ahead order of MGU, by fine-tuning the home positions in the VN and SC pairs.

**Figure 2.**
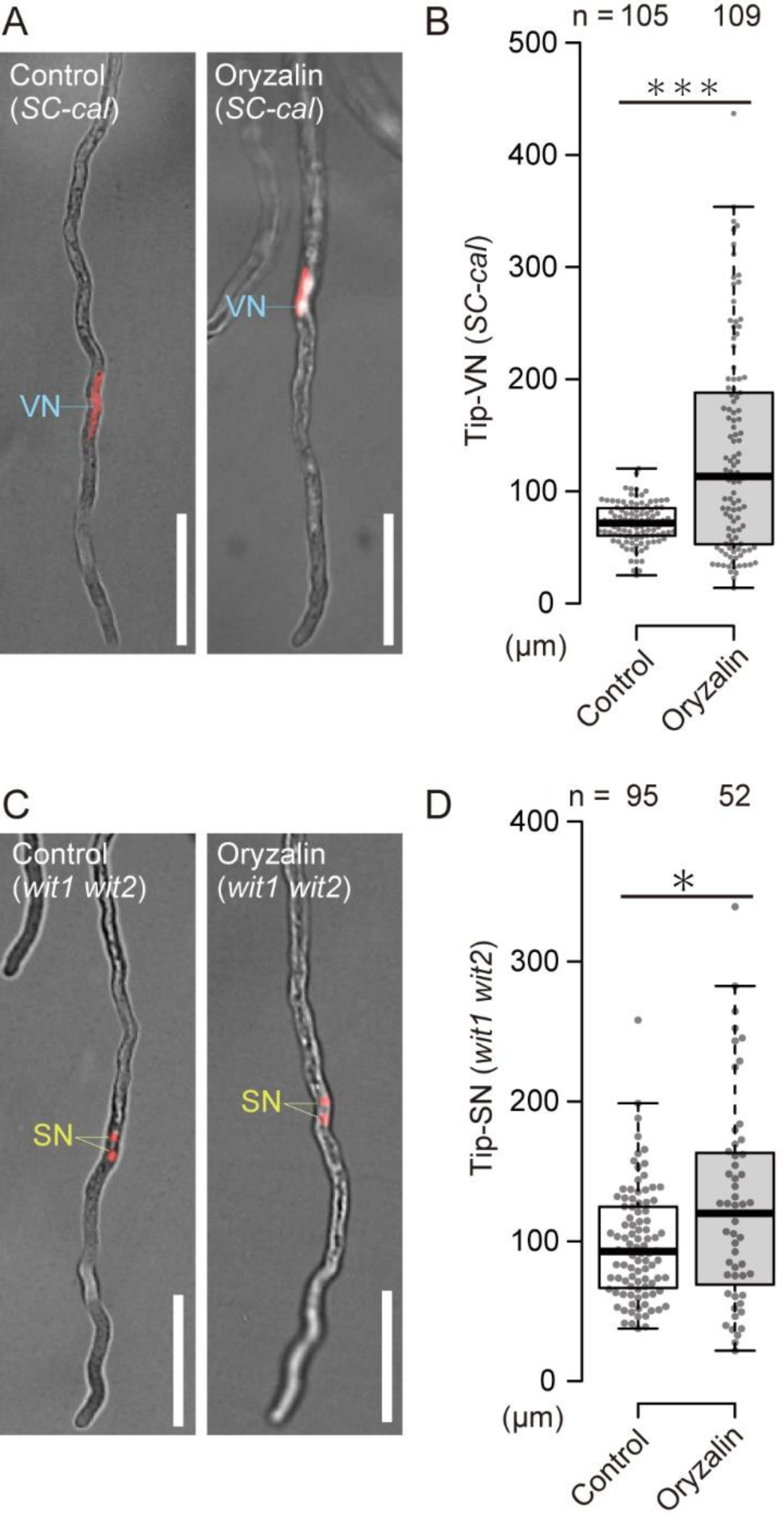
Microtubule-disrupting drug disturbs home positions of the vegetative nucleus and sperm cell pair. **A)** Representative images of an orphan vegetative nucleus under semi-*in vitro*-germinated pollen tubes from the *SC-cal pRPS5A:HISTONE H2B-tdTomato* (*RHT*) plant growing under control or oryzalin-treated conditions. **B)** Distance from pollen tube-tip to vegetative nucleus (Tip-VN) analyzed in (**A**). **C)** Representative images of orphan sperm nuclei from semi-*in vitro*-germinated pollen tubes from the *wit1 wit2 RHT* plant growing under control or oryzalin-treated conditions. **D)** Distance from pollen tube-tip to sperm nuclei (Tip-SN) analyzed in (**C**). In (**A**) and (**C**), red fluorescence signals were observed 6 h after pollination. Box-and-whisker plots show median (center line), upper and lower quartiles (box), maximum and minimum (whiskers), and points (solid circles). Asterisks, significance by Welch’s *t* test (\*\*\**P* = 2.97 × 10^−10^, \**P* = 0.014). Scale bars, 50 μm.

**Figure 3.**
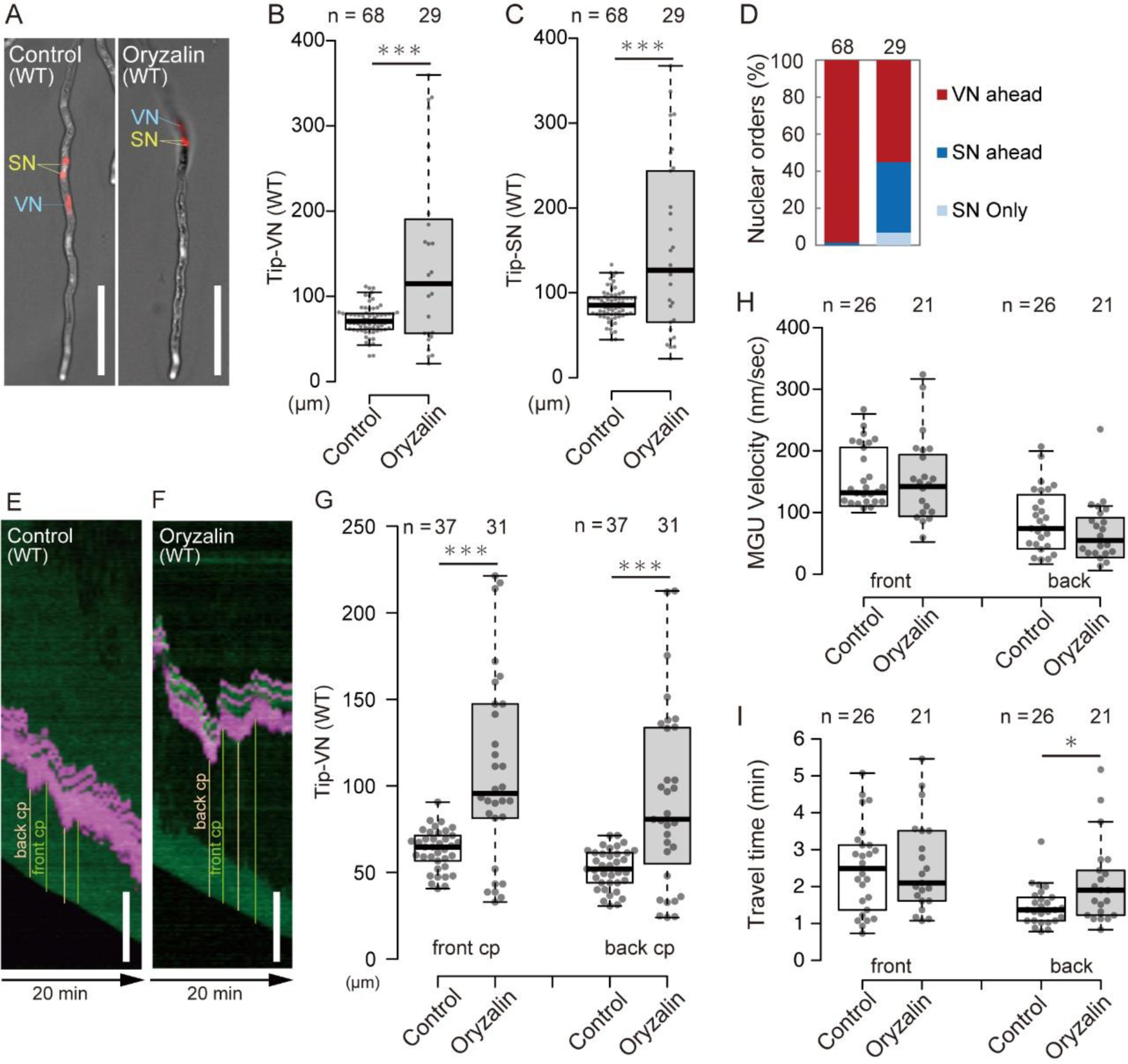
Disruption of male germ unit (MGU) home positions in pollen tubes with disrupted microtubules. **A–D)** Pollen tubes from wild-type (WT) *pRPS5A:HISTONE H2B-tdTomato* (*RHT*) nuclear marker were grown under semi-*in vivo* conditions using control medium or microtubule-unstable medium containing 1 μM oryzalin, and nuclear positions were analyzed 6 h after pollination. Representative images of red fluorescence merged with bright field (**A**), distance from pollen-tube tip to the vegetative nucleus (Tip-VN) (**B**), distance from pollen tube-tip to sperm nuclei (Tip-SN) (**C**), and order of MGU (**D**). **E, F)** Kymographs of MGU movement in pollen tubes on control medium (**E**) or oryzalin-containing medium (**F**). Pollen tubes from WT *RHT pACA3:Lyn24-mNeonGreen* plants were grown under semi-*in vivo* conditions, and time-lapse images were captured every 30 s, 6 h after pollination. **G)** Distance from pollen-tube tip to Tip-VN was measured at the front critical point (cp) or back cp analyzed in (**E**) and (**F**). **H, I)** Velocity (**H**) or travel time (**I**) of MGU migration during forward (front) or backward (back) movement analyzed in (**E**) and (**F**). Box-and-whisker plots show median (center line), upper and lower quartiles (box), maximum and minimum (whiskers), and points (solid circles). Asterisks, significance by Welch’s *t* test (\*\*\**P* < 0.001, \**P* = 0.023). Scale bars, 50 μm.

To determine the features of microtubule-mediated positional control, we performed time-lapse imaging of pollen tubes under semi-*in vivo* growth conditions (Fig. 3E). The movement of MGUs in the control pollen tubes was consistent with other reports; MGUs moved forward toward the apical region as fine saltatory behavior (Schattner et al., 2021). We named the moments when MGUs began to move forward or backward as the front critical point (cp) or back cp, respectively, and measured the distance from the tip to the VN at that moment. In control PTs, both the front cp and back cp showed little variation, suggesting that MGUs accurately recognized their position and maintained their home position by switching between forward and backward movements (Fig. 3, E and G, control). In contrast, under oryzalin treatment, the front cp and back cp values varied greatly, indicating that the precise control of the MGU position was lost (Fig. 3, F and G, Oryzalin). However, there was little difference in the MGU velocity between the control and oryzalin-treated pollen tubes (Fig. 3H). Interestingly, the duration of backward movement was longer in oryzalin-treated pollen tubes than in control pollen tubes (Fig. 3I). These results suggest that microtubules contribute to quick switching between the forward and backward modes of MGUs and consequently control the home position of the MGU.

### *tub4* plants show abnormal MGU positioning and pleiotropic fertilization defects

To obtain genetic evidence of microtubule-mediated MGU positioning, tubulin mutants were analyzed. From a characteristic screwed plant body morphology under microtubule-disrupting conditions, a study identified many semidominant tubulin mutants using a forward genetic approach (Ishida et al., 2007). The mutant collection included plants with missense mutations in four *TUBULIN ALPHA* genes (*TUA2*, *TUA3*, *TUA4*, and *TUA6*) and four *TUBULIN BETA* genes (*TUB1*, *TUB2*, *TUB3*, and *TUB4*). Due to its strong expression in mature pollen, we observed *TUB4* mutants (Loraine et al., 2013). *RHT* was introduced into *tub4^G96D^*, *tub4^L250F^,* and *tub4^S351F^* homozygous mutants, and MGU patterns were compared with those of wild-type *RHT* pollen tubes in *in vitro*-germinated pollen tubes (Fig. 4). In contrast to that observed after oryzalin treatment, we did not find aberrant MGUs whose SC pairs proceeded with the VN (Figs. 3, A and D; 4A). However, *tub4^G96D^*, *tub4^L250F^,* and *tub4^S351F^* homozygous plants showed a broad backward shift in the positions of the VN and SC pairs compared to the wild type (Fig. 4, B and C).

**Figure 4.**
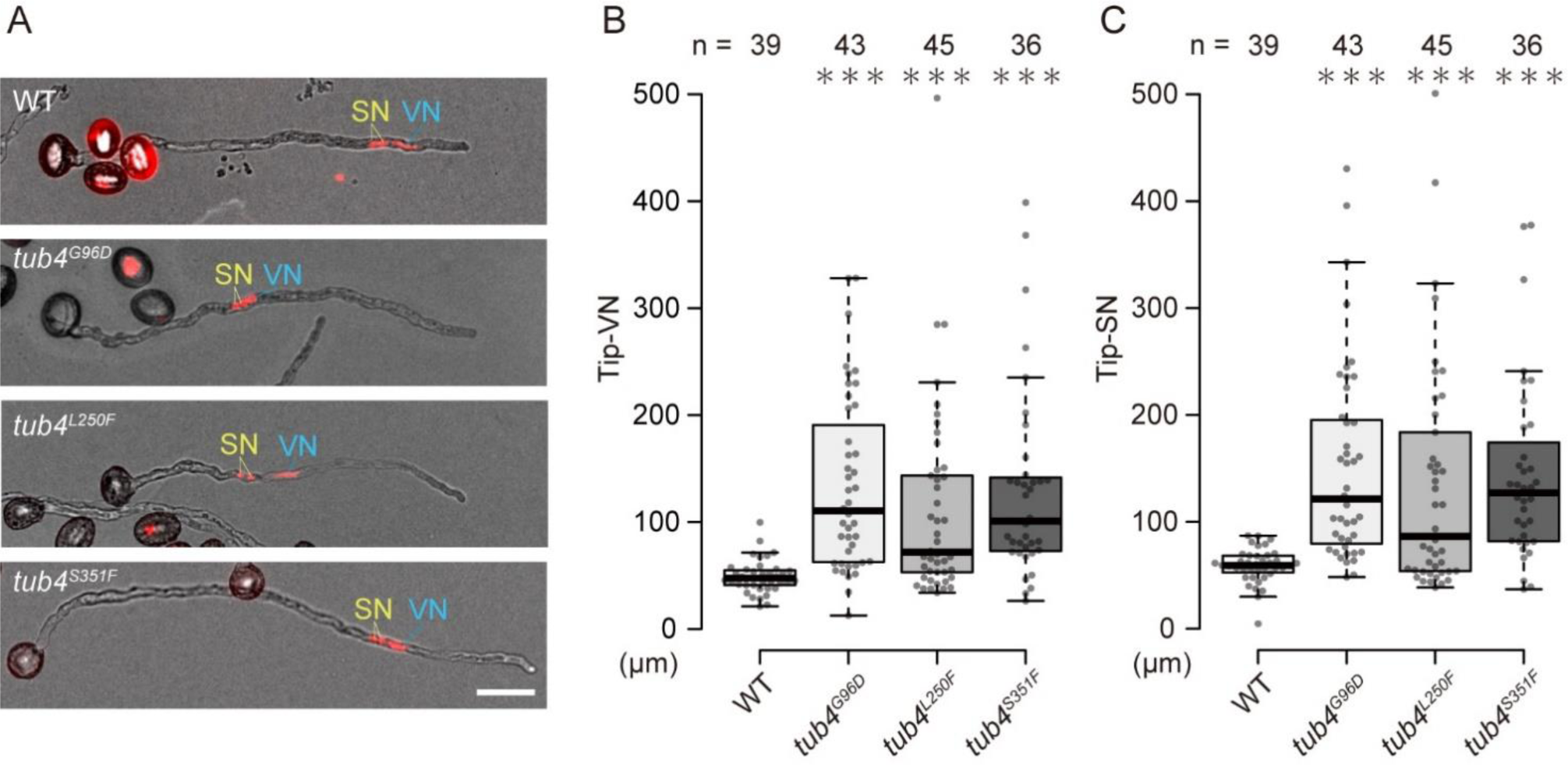
Abnormal male germ unit (MGU) positions in *tub4* plants. **A)** Representative images of *in vitro*-germinated pollen tubes from wild-type (WT) and *tub4* plants carrying the *pRPS5A:HISTONE H2B-tdTomato* (*RHT*) nuclear marker. **B)** Distance from the pollen tube-tip to vegetative nucleus (Tip-VN) analyzed in (**A**). **C)** Distance from the pollen tube-tip to sperm nuclei (Tip-SN) analyzed in (**A**). Red fluorescent signals were observed with transmitting light 6 h after pollination. Box-and-whisker plots show median (center line), upper and lower quartiles (box), maximum and minimum (whiskers), and points (solid circles). Asterisks indicate statistically significant differences detected by Dunnett’s test compared with the WT (\*\*\**P* < 0.001). Scale bars, 50 μm.

To examine the fertility of *tub4* plants, developing siliques from *tub4^G96D^*, *tub4^L250F^*, *tub4^S351F^*homozygous, and wild-type Columbia-0 (Col-0) plants were observed 8 days after manual self-pollination (Supplemental Fig. S2, A and B). Under our growth conditions, wild-type siliques produced normal seeds in approximately 80% of the ovules. In contrast, a moderate reduction in fertility was observed in *tub4^G96D^* (64.7%) and *tub4^L250F^* (63.0%) plants and severely low fertility (13.6%) in *tub4^S351F^* plants (Supplemental Fig. 2B). To distinguish between paternal and maternal fertility defects, we performed reciprocal crosses between wild-type and *tub4* plants (Fig. 5). When *tub4* plants were pollinated with wild-type pollen, all three mutants produced normal seeds comparable to those of wild-type plants (Fig. 5, A and C). However, a significant fertility defect was observed when the wild-type pistils were pollinated with *tub4* pollen (Fig. 5, A and B). The reduction in the normal seed ratio was comparable to the results obtained for autopollination (Fig. 5, A and B; Supplemental Fig. S2B). These results demonstrate male gametophyte-specific dysfunction in *tub4* plants.

**Figure 5.**
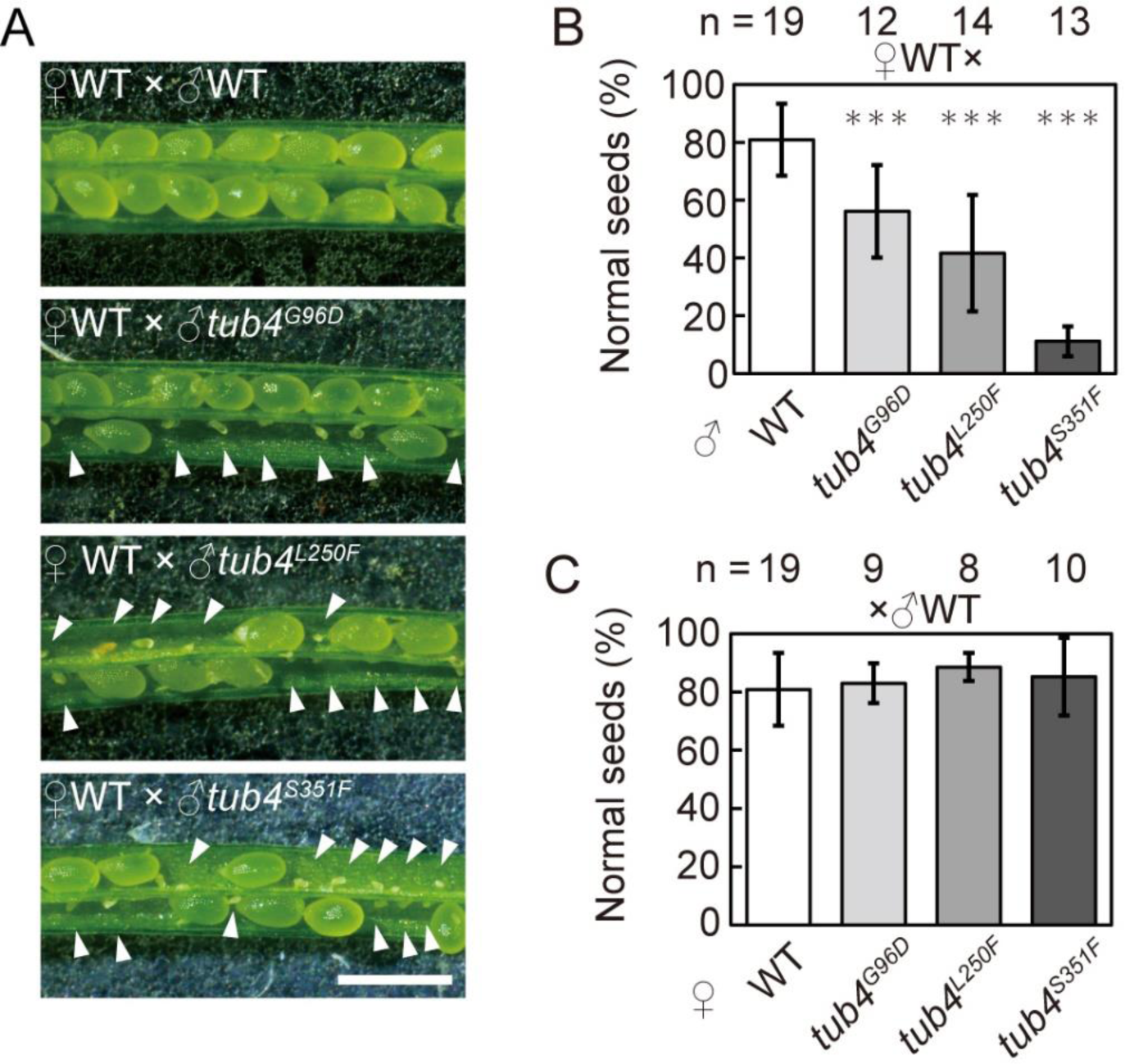
Seed abortion caused by *tub4* pollen tubes. **A)** Representative images of seeds 8 d after wild-type (WT) pistils were pollinated with pollen from WT, *tub4^G96D^*, *tub4^L250F^,* and *tub4^S351F^* plants. **B, C)** Percentages of normal seeds in reciprocal crosses between WT and *tub4* plants. Cross-pollinations using *tub4* pollen donors (**B**). Cross-pollinations using *tub4* plants as female parents (**C**). Scale bar, 1 mm. Arrowheads indicate unfertilized ovules or undeveloped seeds. Asterisks indicate statistically significant differences detected by Dunnett’s test in comparison with the WT (\*\*\**P* < 0.001).

To evaluate reduced fertility in *tub4* plants, we examined pollen-tube growth patterns by aniline blue staining of pistils 24 h after pollination (Supplemental Fig. S3, A to H). In wild-type pistils pollinated with pollen from wild-type plants, the longest pollen tubes reached the bottom of the pistils (Supplemental Fig. S3A, white arrow), and 90.1% of the ovules displayed insertion of pollen tubes (n = 201, Supplemental Fig. S3, B and H). By contrast, *tub4^S351F^*pollen tubes did not reach the bottom of the pistils (Supplemental Fig. S3A). Consequently, only 61.2% of ovules received *tub4^S351F^* pollen tubes (n = 221, Supplemental Fig. S3, E and H). Remarkably, one-half of the ovules in *tub4^S351F^* pistils remained undeveloped even after pollen-tube insertion (comparison of Supplemental Fig. S3, E and H with Fig. 5B). The ovules that did not develop after pollen-tube insertion were distinctly smaller than normally fertilized ovules 48 h after pollination. (Supplemental Fig. S3, I and J, yellow double arrowheads). Failure of double fertilization allows the ovule to attract a second pollen tube and causes multiple pollen-tube insertions, known as polytubey (Hater et al., 2020; Sugi and Maruyama, 2023). Consistently, we observed a slight increase in polytubey in *tub4^S351F^* pollen tubes (Supplemental Fig. S3, F and H). In contrast to *tub4^S351F^*, *tub4^G96D^* and *tub4 ^L250F^* pollen tubes reached the bottom of the pistils (Supplemental Fig. S3A). The frequency of pollen-tube insertion between wild-type and *tub4^G96D^* plants was comparable (84.4%, n = 212, Supplemental Fig. S3, C and H). By contrast, *tub4 ^L250F^* pollen tubes exhibited a slight reduction in ovule insertion (70.5%, n = 268, Supplemental Fig. S3, D and H). In addition, *tub4 ^L250F^*pollen tubes displayed polytubey (4.9%) or overgrowth in the ovular micropyle (5.2%) (Supplemental Fig. S3, F to H). Despite the pleiotropic defects, all three *tub4* mutants displayed a common phenotype of undeveloped seeds produced following insertion of pollen tubes. Most likely, a major fertilization defect in *tub4* plants exists during or after pollen-tube reception.

### Loss of KINESIN-13A caused a forward shift in the home position of MGUs

To examine microtubule function in MGU positioning, we analyzed KINESIN-13 family proteins, which are unique kinesins that lack motor domains and induce microtubule depolymerization (Oda and Fukuda, 2013). The expression of *KINESIN-13* in pollen tubes was reported in early immunostaining (Wei et al., 2005). Among the two *KINESIN-13* (*KIN13*) genes encoded in the *Arabidopsis* genome, only *KIN13A* is expressed in the pollen and pollen tubes (Qin et al., 2014). Thus, pollen-tube growth and MGU positions were analyzed in wild-type, *kin13a*, and two complementation lines (*kin13a*/*eYFP KIN13A*) *in vitro* using medium containing the DNA fluorescent dye Kakshine532 (Uno et al., 2021). *kin13* pollen tubes displayed normal MGU order and grew to the same length as wild-type pollen tubes (Fig. 6, A and B). However, the VN of *kin13* pollen tubes was frequently detected at more tip-close positions than that in wild-type (Fig. 6, A and C). The forward shift of VN positioning was rescued in the two complementation lines (Fig. 6, A and C). These data suggest that KIN13A is a microtubule depolymerizing agent that controls the tip-to-MGU distance.

**Figure 6.**
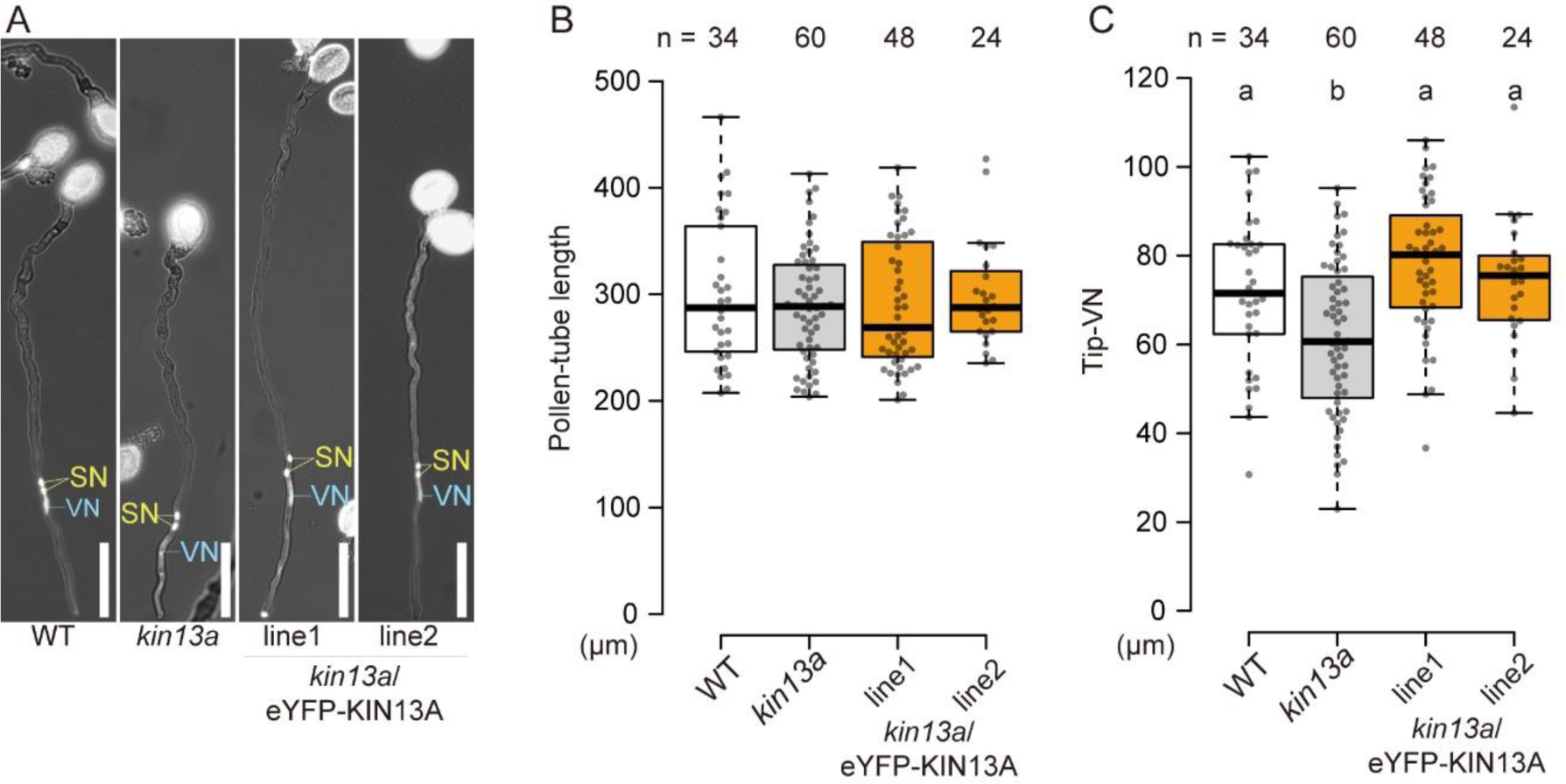
Kinesin-13A modulates male germ unit (MGU) position. **A)** Representative images of pollen tubes from wild-type (WT), *kin13a*, *kin13a eYFP-KIN13A* plants. Pollen tubes were *in vitro*-germinated on growth medium containing 1 μM Kakshine532 fluorescent DNA dye and analyzed after 2.5 h incubation. Scale bar, 50 μm. **B, C)** Pollen tube length (**B**) and distance from pollen tube-tip to vegetative nucleus (Tip-VN) (**C**) analyzed in (**A**). Box-and-whisker plots show median (center line), upper and lower quartiles (box), maximum and minimum (whiskers), and points (solid circles). Different alphabets indicate statistically significant differences detected by Tukey’s test.

### Observations of tubulin reporters in pollen tubes

Inhibitor and mutant analyses suggested the significance of spatiotemporal microtubule regulation in MGU positioning. We introduced translational sGFP fusion reporter genes of *TUA1*, *TUA2*, *TUA6*, or *TUB4* into *RHT* and performed time-lapse imaging of microtubular patterns under semi-*in vivo* growth conditions (Fig. 7). These reporter lines exhibited a bright GFP signal in the vegetative cell of the pollen tube. Because the GFP signal in the vegetative cell, the microtubular pattern could not be clearly determined in the SCs of *pTUA1:sGFP-TUA1*, *pTUA2:sGFP-TUA2*, and *pTUB4:sGFP-TUB4* lines (Fig. 7). Although the *pTUA6:sGFP-TUA6* lines only showed a strong GFP signal in the SCs, microtubular patterns were barely detectable (Fig. 7). In contrast, several common microtubular patterns were visible in the vegetative cells of the growing pollen tube (Fig. 7; Supplemental Videos 1 to 4). First, sGFP-labeled tubulin formed a moderate gradient from pollen-tube tip to the basal region. Second, actively moving short microtubules accumulated in front of the MGU, whereas longer cables were distributed around the basal region of the MGU. Third, longer static cables were found around the periphery of the plasma membrane. Fourth, rapid microtubule movements were observed at the center of pollen tubes, which sometimes appeared to associate with VN or SC pairs (Fig. 7; Supplemental Videos 1 to 4). Unfortunately, quantitative analysis of the association between microtubules and each MGU component was hindered by vigorous protoplasmic streaming.

**Figure 7.**
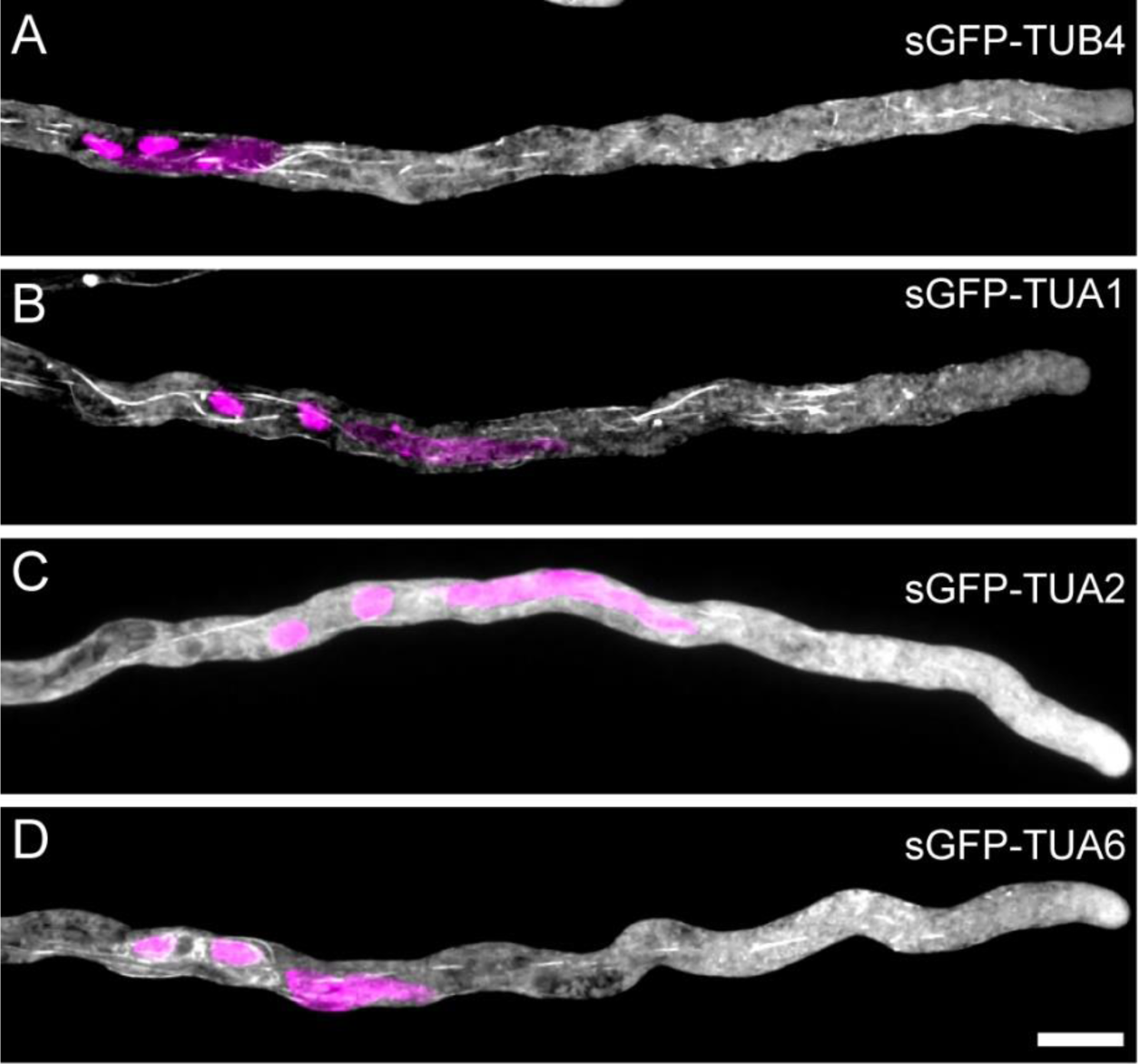
Microtubule dynamics during male germ unit (MGU) transport. **A–D)** Microtubules (gray) and nuclei (magenta) in pollen tubes from *pRPS5A:HISTONE H2B-tdTomato* (*RHT*) *pTUB4:sGFP-TUB4* (**A**), *pTUA1:sGFP-TUA1* (**B**), *pTUA2:sGFP-TUA2* (**C**), or *pTUA6:sGFP-TUA6* (**D**) plants. Pollen tubes were grown under semi-*in vivo* conditions at 6 h after pollination. Scale bar, 10 μm.

Time-lapse imaging showed clear abnormalities in GFP patterns or pollen-tube behaviors in the tubulin reporter lines. For example, the *pTUB4:sGFP-TUB4* line showed severe pollen-tube growth retardation and backward shift of MGU, although visible microtubules were most frequently observed in the four tubulin reporters (Fig. 7; Supplemental Fig. S4). Similar but moderate defects were detected in the other reporter lines. The *pTUA1:sGFP-TUA1* lines often produced granule-shaped structures, in addition to cable patterns, indicating the formation of nonnative protein aggregates. These observations suggest formation of artifacts due to the overexpression of sGFP-tagged tubulin proteins. The number of visible microtubule fragments decreased during time-lapse imaging. This alteration would be partially due to photobleaching, but likely shows higher photosensitivity of microtubule dynamics in pollen tubes. Taken together, the artifacts of tubulin reporter lines, higher photosensitivity, and rapid protoplasmic streaming demonstrate challenges in analyzing the native dynamics of microtubules in pollen tubes.

## Discussion

Apical MGU transport is a key mechanism of pollen tube-dependent sexual reproduction in flowering plants. In this study, the genetic dissection of MGU motility indicated that VN and the SC pair tended to be located at different positions in the apical region of the pollen tube, and MGU positioning was disturbed in *tub4* and *kin13a* plants. These findings suggest independent home positions of VN and the SC pair produced through unique saltatory movements modulated by microtubule dynamics.

### Live imaging of tubulin in pollen tubes

According to antitubulin immunostaining studies in *Nicotiana tabacum*, *Nicotiana alata, Pyrus communis*, and *A. thaliana*, pollen tubes show characteristic microtubular patterns: the tip region contains no microtubules, the subapical region in front of the MGU displays a distribution of shorter bundles, and the shank region around and beneath the MGU produces longitudinally aligned cortical microtubules (Heslop-Harrison et al., 1988; Tiwari and Polito, 1988; Del Casino et al., 1993; Joos et al., 1994; Åström et al., 1995; Joos et al., 1995; Raudaskoski et al., 2001; Laitiainen et al., 2002; Yu et al., 2009). In addition to fixed sample observations, microtubule movements were reported in tobacco pollen tubes transiently expressing GFP-AtEB1a through microprojectile bombardment (Cheung et al., 2008). During live imaging, pollen tubes showed a dense signal of GFP-AtEB1a in the subapical region and differential microtubular dynamics, including static cortical microtubules in the shank and rapid flux of short microtubule bundles in the subapical region (Cheung et al., 2008). Our time-lapse imaging of four sGFP-tagged tubulin reporters confirmed these observations (Fig. 7, Supplemental Videos 1 to 4). However, direct GFP-tagging exhibits a moderate gradient of tubulins from pollen-tube tip to the basal region and implies a different spatiotemporal control of the polymerization–depolymerization equilibrium along the longitudinal axis of the pollen tube.

### Microtubule-mediated MGU transport and the tug-of-war model

A study proposed the association of microtubules with MGU using immunostaining and electron microscopy data (Lancelle et al., 1987). Bright field analysis of growing pollen tube revealed a dynamic association between MGU and fibrillar structures (Heslop-Harrison and Heslop-Harrison, 1989b). Our live imaging confirmed the close presence of microtubules around the MGU (Fig. 7; Supplemental Videos 1 to 4). Microtubule disruption by oryzalin or colchicine treatment retarded the apical transport of VN and generative cells in *N. tabacum*, *N. alata, N. sylvestris*, and *Galanthus nivarius* (Heslop-Harrison et al., 1988; Joos et al., 1994; Joos et al., 1995; Åström et al. 1995; Laitiainen et al., 2002). A similar broad MGU distribution phenotype was observed in oryzalin-treated *Arabidopsis* pollen tubes (Fig. 3), indicative of apical MGU transport regulated by microtubules. By contrast, cytochalasin D-mediated F-actin disruption had a more deleterious effect on apical MGU transport (Heslop-Harrison and Heslop-Harrison, 1989a, 1989b). Thus, microtubules play a subsidiary role in apical MGU transport. Recently, Schattner et al. proposed a tug-of-war model, in which KCH kinesins drive bimodal SC behavior, consisting of rapid forward and slow backward movements (Schattner et al., 2021). The tug-of-war model largely owes to KCH activity, which generates different velocities from the parallel or antiparallel sliding of F-actin and microtubules (Walter et al., 2015), suggesting a key role of microtubular dynamics during MGU transport. A microtubule cage, which is a longitudinally aligned microtubule bundle in generative cells and SCs, is the primary candidate of a KCH-associated microtubule scaffold (Schattner et al., 2021). Although microtubule cages are resistant to oryzalin in several species (Heslop-Harrison et al., 1988; Åström et al. 1995; Zhang et al., 1995; Laitiainen et al., 2002), *Arabidopsis* pollen tubes appeared to lack microtubule cages, irrespective of oryzalin treatment, based on observations of antitubulin immunostaining and microtubule reporters, including *pUBQ14:GFP-TUA6* and other tubulin markers in this study (Fig. 7) (Yu et al., 2009; Oh et al., 2010). Electron microscopy analysis revealed that a microtubule cage is an assembly of cortical microtubules inside generative cells or SCs and is unable to participate in KCH-mediated cable sliding expected in the tug-of-war model (Lancelle et al., 1987). Finally, our study demonstrated that in *Arabidopsis*, oryzalin did not affect MGU velocity during forward or backward movement (Fig. 3H). These data suggest amicrotubule-independent bimodal MGU movement. Overall, the major generators of bimodal MGU movement are probably F-actin and myosins, and not a microtubule-dependent system, as argued in other studies (Cai and Cresti, 2010).

### Role of microtubules in MGU positioning

Our time-lapse imaging recorded a longer backward traveling time of MGU after oryzalin treatment (Fig. 3I). Microtubules support stable apical MGU positioning presumably by modulating the timing of direction switching during basipetal MGU movement. This MGU positioning control probably requires the homeostasis of microtubular patterns dependent on the equilibrium between tubulin polymerization and depolymerization. Compromised MGU positioning due to oryzalin treatment or *tub4* mutations should be a representative phenotype of microtubule instability (Figs. 2 to 4). Interestingly, our sGFP-tagged tubulin reporter lines showed broad MGU positioning (Fig. 7; Supplemental Fig. S4; Supplemental Video 1). These reporter lines have higher copies of the tubulin gene, which is expected to increase tubulin polymerization, in contrast to oryzalin treatment. Yu et al. performed antitubulin immunostaining in *Arabidopsis* pollen expressing *TUA1* from *Picea wilsonii* under a strong *LAT52* promoter. Compared with the wild-type pollen tubes, the transgenic pollen tubes enhanced the α tubulin signal and increased the accumulation of short microtubules in the apical-to-subapical region (Yu et al., 2007). Most likely, tubulin overexpression extends the subapical region occupied with short microtubules in front of the MGU, causing MGU positioning to shift backward.

Because pollen tubes penetrate with vigorous protoplasmic streaming, they must have a sophisticated system that maintains a unique equilibrium of microtubule polymerization–depolymerization in each region. KIN13A can be a component of this homeostasis. In *Arabidopsis* mesophyll protoplasts, *At*KIN13A is present in the Golgi apparatus (Lu et al., 2005). This subcellular localization was confirmed in pollen tubes of *N. tabacum*, the KIN13A ortholog (*Nt*KIN13As) of which was detected by anti-*At*KIN13A immunostaining (Wei et al., 2005). Thus, KIN13A was speculated to be involved in protein secretion at the pollen-tube apex (Wei et al., 2005; Cai and Cresti, 2010). Analysis of metaxylem vessel cells in *Arabidopsis* showed that KIN13A plays a pivotal role in secondary cell wall pit formation by inducing depolymerization of cortical microtubules necessary for sliding movement of the cellulose synthase complex (Oda and Fukuda, 2013). KIN13A is recruited to the cortical microtubules by an active form of Rho of plants (ROP) GTPase and MICROTUBULE DEPLETION DOMAIN1 (MIDD1) protein. In pollen tubes, ROP is localized at the apex, where it controls polar protein secretion required for active tip growth (Hwang and Yang, 2006). In this study, *kin13a* pollen tubes showed a forward shift of MGU positioning (Fig. 6). It is possible that KIN13A destroys cortical microtubules at the apical region downstream of ROP signaling, and this local microtubule instability maintains the size of the subapical region with short microtubule bundles. Although microtubules do not play a major role in pollen-tube growth (Joos et al., 1994; Cai and Cresti, 2010), two microtubule-destabilizing proteins (MICROTUBULE-ASSOCIATED PROTEIN18 [MAP18] and plasma membrane-associated cation binding protein 1 [PCaP1]/MICROTUBULE-DESTABILIZING PROTEIN25 [MDP25]) have been shown to control the directional growth of pollen tubes by severing F-actin at the subapical region (Wang et al., 2007; Nagasaki et al., 2008; Kato et al., 2010; Li et al., 2011; Zhu et al., 2013; Qin et al., 2014). These microtubule-destabilizing proteins may control MGU positioning in parallel with the KIN13A pathway.

### Independent control of home positions in VN and the SC pair

Upon pollen tube discharge, the pollen tube must include two SCs in a limited volume of released cellular contents. If MGU positioning is shifted backward, the pollen tube cannot deliver SCs and fails to accomplish double fertilization. Pollen tubes from *tub4* plants showed reduced fertility even after ovule arrival (Figs. 4 and 5). Pollen tubes from *SC-cal* plants exhibited reduced fertility due to abnormal SC positioning (Motomura et al., 2021). Our analysis of *tub4* plants provides additional evidence on the crucial role of MGU positioning in double fertilization.

Genetic dissection of MGU motility using *SC-cal* and *wit1 wit2* plants revealed independence of home positions between VN and the SC pair (Fig. 1). Due to their own home positions, MGUs could maintain the VN-ahead order during transport (Fig. 3D). Although the VN-ahead MGU order is common in flowering plants (Heslop-Harrison and Heslop-Harrison, 1989c), its biological significance is unclear. SC pairs or generative cells may receive less mechanical stress than the opposite order, protected by the proceeding VN. Concomitantly with a robust VN–SC connection, the VN-ahead MGU order and different home positions in MGU would ensure stable male genome delivery.

Compared with other pollen-tube biology studies, studies of MGU transport have progressed slowly, probably due to the tight association of F-actin function in pollen-tube growth and technical difficulty in dissecting bimodal movement. Our data might change this situation by focusing on microtubule function and proposing the home position concept. Discovery of independent home positions indicated putative molecular switches on VN and SCs that perceive relative distance from the tip and determine the timing to change moving directions. Candidates for switch proteins can be selected from proteomics data in MGU. To characterize new positioning regulators, monitoring the native microtubular pattern is necessary. Because the increased copy number of tubulin in our reporter lines induced an artificial microtubular pattern, more authentic reporter lines should be generated using in-frame GFP knock-in technology (Miki et al., 2018). These analyses of the MGU positioning system will contribute to the genetic dissection of basic bimodal MGU movement in the future.

## Methods

### Plant materials and growth conditions

Col-0 was used as the background for all plants. *RHT*, *SC-cal RHT*, *wit1 wit2 RHT*, and *Lyn24-mNeongreen RHT* plants have been described (Motomura et al., 2021). *tub4^G96D^* (CS68892), *tub4^L250F^* (CS68896), and *tub4^S351F^* (CS68897) seeds were provided by ABRC (https://abrc.osu.edu). *kin13a* and *kin13a*/*eYFP-KIN13a* plants were gifts from Dr. Yoshihisa Oda (Nagoya University). *ms1* plants were generated using CRISPR/Cas9-mediated genome editing. Sequencing showed insertion of a G at 1,166 bp from the first base of the start codon, A, leading to a frameshift. After confirming Cas9-null status in the T2 generation, T3 *ms1* homozygous mutants were selected based on phenotype, and the pistils of these plants were used for semi-*in vitro* pollen-tube growth.

Seeds were directly sown on soil or were surface sterilized with sterilization solution (2% Plant Preservative Mixture, 0.1% Tween 20), incubated at 4°C for several days, and sown on Murashige and Skoog medium containing 1% sucrose and antibiotics. Approximately 10 d after germination, the plants were transferred to the soil and grown at 22°C under continuous light or standard long-day conditions.

### Plasmids and transgenic plants

Plasmids carrying sGFP-tubulin reporter genes were constructed as follows: In each tubulin gene, the promoter sequence and protein coding sequence with 3′-UTR were independently amplified by PCR using primer sets listed in Supplemental Table S1 and Col-0 genome DNA as a template. DNA fragment encoding sGFP with the Gly-Gly-Ser-Gly-Gly-Ser linker was amplified by PCR using mNG_F and GFP_Linker_R as primer sets from the pDM310 plasmid containing *pMYB98:pCOXIV-sGFP* (Maruyama et al., 2015). DNA cassettes of the promoter, protein coding sequence, and sGFP were mixed and subjected to PCR using the forward primer used for the amplification of the promoter and the reverse primer used for the amplification of the protein coding sequence. These PCR products were introduced into pDONR221 by BP reactions to produce four sGFP-tubulin entry vectors (Thermo Fisher, MA, USA). We performed the LR reaction of the pGWB501 destination vector with sGFP-tubulin entry vectors carrying *pTUA1:sGFP-TUA1*, *pTUA2:sGFP-TUA2*, *pTUA6:sGFP-TUA6*, or *pTUB4:sGFP-TUB4*, and obtained binary vectors pDM772, pDM693, pDM694, and pDM774, respectively (Nakagawa et al., 2007). The plasmids were transformed into *RHT* plants using the *Agrobacterium*-mediated floral dipping method, with *Agrobacterium* GV3101 (Zhang et al., 2006). After kanamycin selection, T_2_ homozygous plants were used for live imaging.

### *In vitro* pollen-tube growth

Mature pollen was collected and observed on pollen germination medium (PGM: 0.01% boric acid, 5 mM CaCl_2_, 5 mM KCl, 1 mM MgSO_4_, 10% sucrose; pH adjusted to 7.5 with 1 N KOH) and 1.5% NuSieve GTG agarose supplemented with 10 μM epibrassinolide) (Muro et al., 2018). Pollen tubes were observed using an upright microscope (Axio Imager. M1; Zeiss, Oberkochen, Germany). To label DNA in sperm nuclei and VN, we used PGM containing 1 μM Kakshine532 (Uno et al., 2021). Fiji (ImageJ image processing package) was used for image processing (Schindelin et al., 2012).

### Semi-*in vitro* pollen-tube growth and oryzalin treatment

Stigmas from *ms1* or emasculated Col-0 pistils were cut with a needle and placed on the medium, and hand pollinated with pollen from transgenic plants. After 6 h of incubation, pollen tubes that emerged from the cut pistils were observed using an Axioscope 5 upright microscope (ZEISS, Oberkochen, Germany) with a CCD camera (Axiocam 503 mono; ZEISS) or an IX73 inverted microscope (Evident, Tokyo, Japan), equipped with a 60× objective lens, a spinning disk confocal scanning unit (Yokogawa Electric, Musashino, Japan), and an sCMOS camera (Zyla 4.2; Andor, Belfast, Northern Ireland). For oryzalin treatment, PGM was thawed at 65°C and cooled to 37°C before adding 0.1% 1 mM oryzalin (final concentration 1 μM) or 0.1% ethanol as control and spread thinly on a dish. Live imaging experiments were performed with an IX83 inverted microscope (Evident) equipped with a confocal scanner unit (CSU-W1; Yokogawa Electric, Musashino, Japan), an sCMOS camera (Zyla 4.2 PLUS; Oxford Instruments, Abingdon, UK), a laser diode illuminator (LDI-7; 89 NORTH, VT, USA), and a silicone immersion objective lens (UPLSAPO100XS2, Evident, Tokyo, Japan). Stack images were acquired at 1-mm increments and 20-s time interval and processed with deconvolution using cellSens Dimension and Maximum Intensity Z-projection in Fiji (Schindelin et al., 2012).

### Seed viability and reciprocal cross

For self-pollination, mutant pistils emasculated a day before were pollinated with pollen from the same mutant. Pollen from the same mutant was pollinated with each mutant pistil. For reciprocal cross, Col-0 or mutant pistils emasculated a day before were pollinated with pollen from Col-0 or mutant anthers. After 8 days, siliques were dissected and seed viability was determined. Images were captured using MVX10 (Evident, Tokyo, Japan) under bright field.

### Aniline blue staining

Pollen was pollinated on Col-0 pistils emasculated a day before. After 24 or 48 h of pollination, pistils were fixed with a fixative (ethanol: acetic acid = 9:1) for 3 h. Fixative was replaced with 1 N sodium hydroxide solution by standing overnight. Samples were immersed in aniline blue staining solution (0.1% aniline blue, 0.1 M K_3_PO_4_) for 10 min. Images were acquired using an IX73 inverted microscope (Evident). Fluorescence signals were detected with a CFP filter using a mercury vapor lamp with a 10× objective lens to observe pollen-tube growth and excited with a 405 nm laser with a 20× or 40× objective lens to observe pollen-tube attraction to the ovule.

## Author Contributions

KM and DM designed the study, conducted the experiments, analyzed pollen-tube growth, and drafted the manuscript. HT performed all analyses of tubulin mutants under the support of NS and DS. MK performed the experiments using oryzalin. AM performed various quantitative analyses. AT constructed a plasmid for genome editing. TK provided critical advice and reviewed the manuscript. All authors have contributed to the manuscript and approved the final version.

## Acknowledgements

We thank K. Tamura and I. Mayer for providing the *wit1 wit2* mutant; D. Kurihara for providing *pRPS5A:H2BtdTomato*; Y. Oda for sharing *kin13a* and *kin13a*/*eYFP-KIN13a* seeds; Y. Sato for providing Kakshine532; A. Saito, Y. Kibayashi, S. Yang, and H. Kakizaki for their technical support. We would like to thank enago for English editing.

