## Supplementary Figures and Table for "Microtubules ensure transport of vegetative nuclei and sperm cells by fine-tuning their home positions"

**Supplemental Figures**

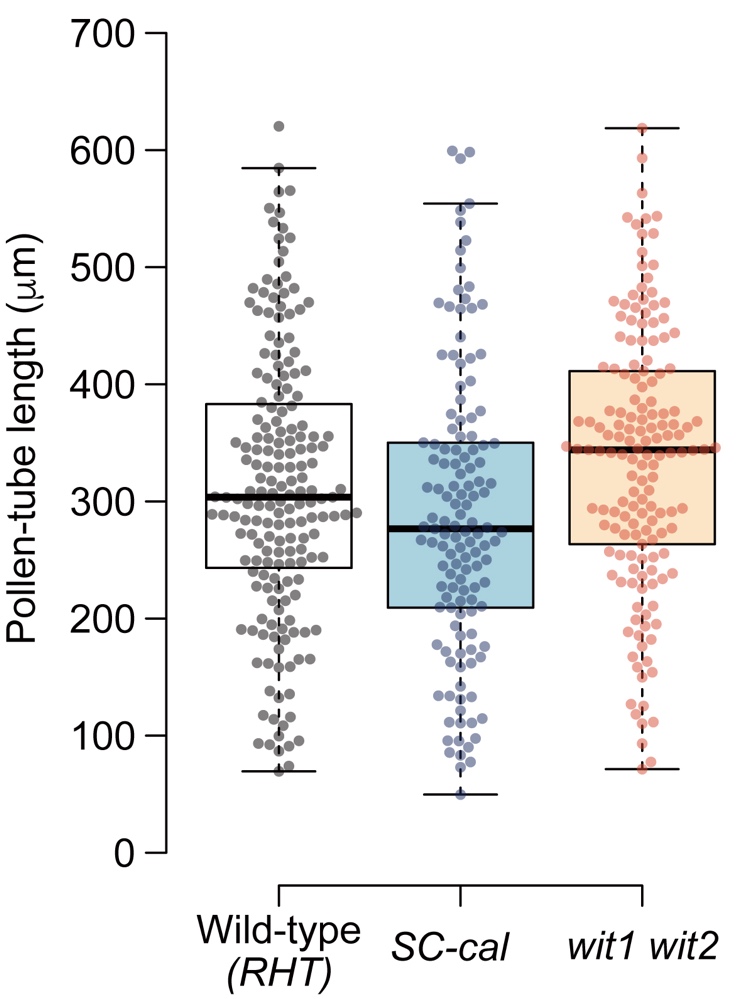

**Supplemental Figure S1. Length of *in vitro*-germinated pollen tubes.**

Pollen from wild-type, *SC-cal* transgenic, and *wit1 wit2* plants carrying the *pRPS5A:HISTONE H2B-tdTomato* (*RHT*) nuclear marker were germinated on pollen-tube growth medium. Length of *in vitro*-germinated pollen tubes were measured 3 h after germination. Box-and-whisker plots show median (center line), upper and lower quartiles (box), maximum and minimum (whiskers), and points (solid circles).

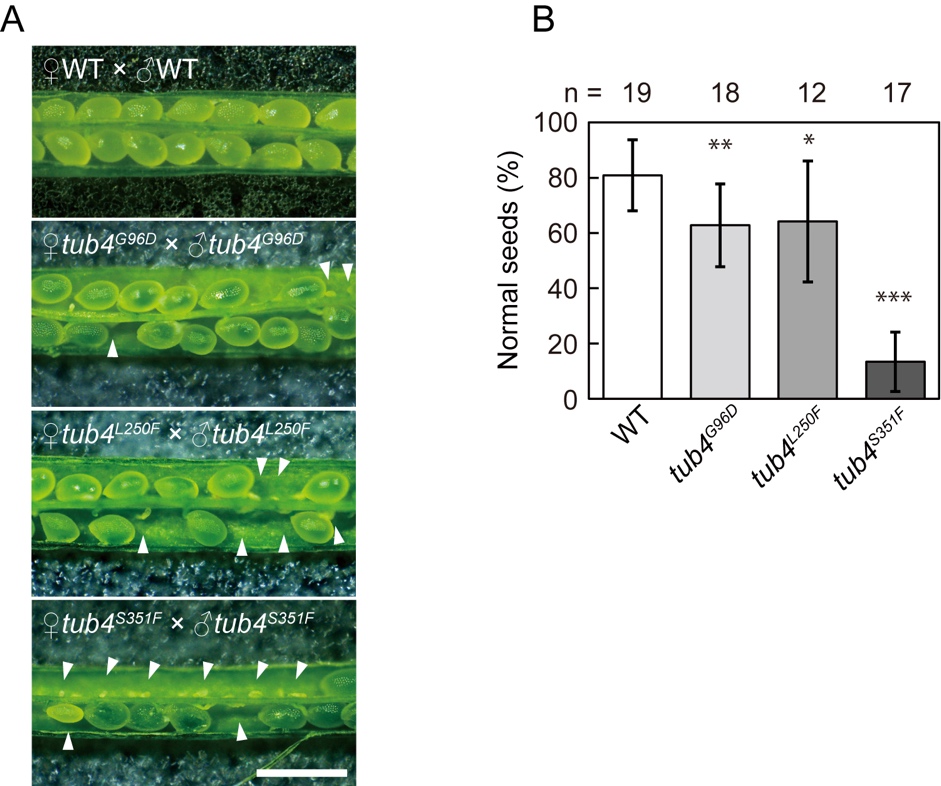

**Supplemental Figure S2. Seed development in *tub4* mutants.**

**A)** Representative images of seeds 8 d after manual self-pollination in *tub4^G96D^*, *tub4^L250F^*, *tub4^S351F^* homozygous, and wild-type Columbia-0 (Col-0) plants. Scale bar, 1 mm. **B)** Percentages of normal seeds in manually self-pollinated wild-type and *tub4* plants. Arrowheads indicate unfertilized ovules or undeveloped seeds. Asterisks indicate statistically significant differences detected by Dunnett’s test compared with WT (**P* = 0.0103, ***P* = 0.0014, ****P <* 0.001).

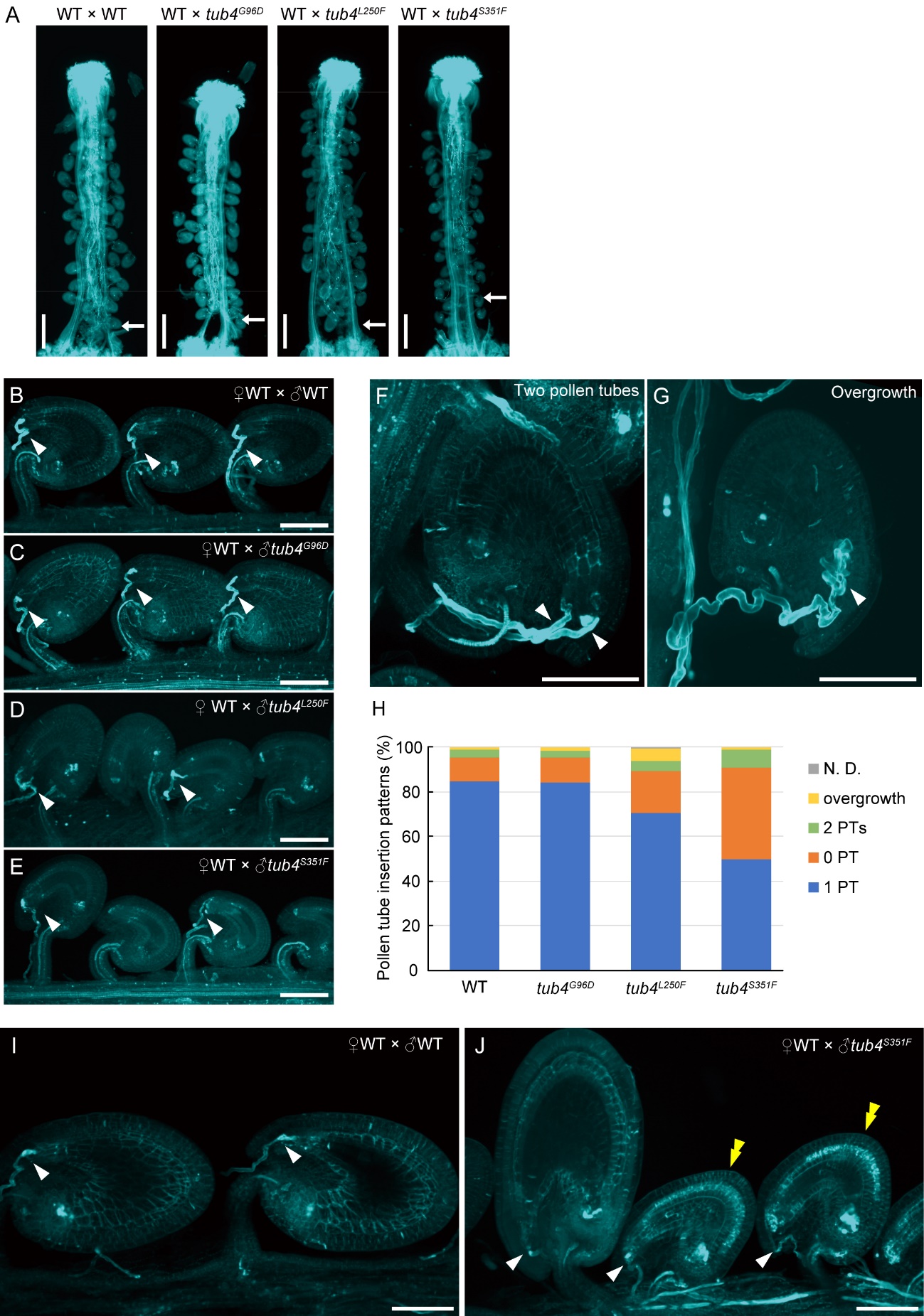

**Supplemental Figure S3.** **Analysis of *in vivo* pollen-tube growth and micropylar guidance in wild-type and *tub4* pollen tubes.**

**A)** Wild-type (WT) pistils were pollinated with pollen from WT, *tub4^G96D^*, *tub4^L250F^,* and *tub4^S351F^* plants, and pollen-tube growth patterns were observed 24 h after pollination by aniline blue staining. Scale bars, 400 μm. **B–E)** Pollen tubes around the micropyle were observed in samples prepared as in (**A**). Scale bars, 100 μm. **F)** WT ovules receiving two pollen tubes. This representative image was captured during observation in (**D**). **G)** WT ovules exhibiting pollen-tube overgrowth. This representative image was captured during observation in (**E**). **H)** Percentages of pollen-tube insertion patterns analyzed in (**B–E**). Ovules receiving no pollen tube, one pollen tube, and two pollen tubes are indicated as 0 PT, 1 PT, and 2 PTs, respectively. N.D.: not determined. **I, J)** Ovules 48 h after pollination. WT pistils were pollinated with pollen from WT (**I**) or *tub4^S351F^* (**J**) plants and observed 48 h after pollination with aniline blue staining. Scale bars, 100 μm. Arrows in (**A**) indicate pollen tubes receiving ovules that are most distant from the stigma. Arrowheads in (**B**) to (**G**), (**I**), and (**J**) show pollen tubes inserted into ovules. Yellow double arrowheads in (**J**) indicate undeveloped ovules after pollen-tube insertions.

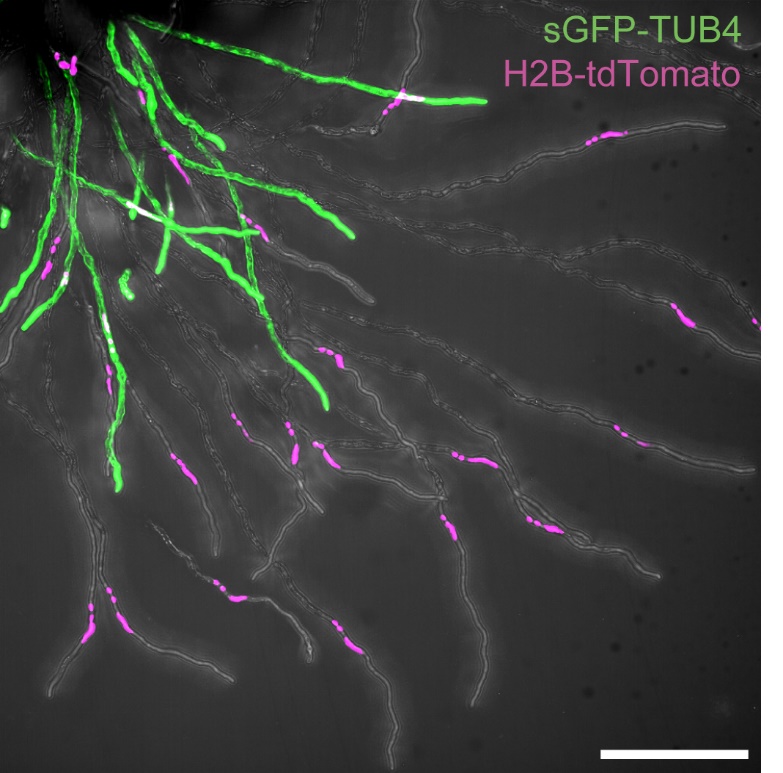

**Supplemental Figure S4. Growth retardation in pollen tubes expressing *sGFP-TUB4*.**

Pollen from a transgenic line homozygous for the *pRPS5A:HISTONE H2B-tdTomato* (*RHT*) nuclear marker and hemizygous for *pTUB4:sGFP-TUB4* were germinated under semi-*in vivo* growth condition, and their pollen tubes were observed 6 h after pollination. GFP-positive pollen tubes often showed retarded growth and a backward shift of male germ unit (MGU), compared with GFP-negative pollen tubes segregated from the same transgenic plant. Scale bar, 100 μm.

**Captions for supplemental Videos**

**Video S1 Representative confocal movie of pollen tubes carrying *pTUB4:sGFP-TUB4* and *pRPS5A:H2B-tdTomato* nuclear marker.** Pollen tubes were grown under semi-*in vivo* conditions. Time stamp, mm:ss.

**Video S2 Representative confocal movie of pollen tubes carrying *pTUA1:sGFP-TUA1* and *pRPS5A:H2B-tdTomato* nuclear marker.** Pollen tubes were grown under semi-*in vivo* conditions. Time stamp, mm:ss.

**Video S3 Representative confocal movie of pollen tubes carrying *pTUA2:sGFP-TUA2* and *pRPS5A:H2B-tdTomato* nuclear marker.** Pollen tubes were grown under semi-*in vivo* conditions. Time stamp, mm:ss.

**Video S4 Representative confocal movie of pollen tubes carrying *pTUA6:sGFP-TUA6* and *pRPS5A:H2B-tdTomato* nuclear marker.** Pollen tubes were grown under semi-*in vivo* conditions. Time stamp, mm:ss.

**Supplemental Table**

| **Table S1 Primers used in this study.** | | |
| --- | --- | --- |
| Amplified cassette | Primer name | Sequence (5′–3′) |
| *TUA1* promoter | TUA1pro_attB1_F | GGGGACAAGTTTGTACAAAAAAGCAGGCTCCCTCCCCCTTCTTTTTTTC |
|  | TUA1pro_XFP_R | CTCGCCCTTGCTCACCATGTTCGAAAATTTCTCAGAGAC |
| *TUA1* protein coding & terminator | Linker1-TUA1_F | GGTGGCAGCGGCGGCAGCATGAGGGAGATCATTAGCATTC |
|  | TUA1_attB2_R | GGGGACCACTTTGTACAAGAAAGCTGGGTTGGTCGAAGATTGTGAAAGACAGGTC |
| *TUA2* promoter | TUA2pro_attB1_F | GGGGACAAGTTTGTACAAAAAAGCAGGCTCGCCTTCTGGTCCAACGATT |
|  | TUA2pro_XFP_R | CTCGCCCTTGCTCACCATTTTCGTTTCGCGGAAGAAAGAAAAGATC |
| *TUA2* protein coding & terminator | Linker1_TUA2coding_F | GGTGGCAGCGGCGGCAGCATGAGAGAGTGCATTTCGATCCACATTG |
|  | TUA2term_attB2_R | GGGGACCACTTTGTACAAGAAAGCTGGGTTGTTCCAAAAATTTCAATCTTTCACCACTCAAATAAC |
| *TUA6* promoter | TUA6pro_attB1_F | GGGGACAAGTTTGTACAAAAAAGCAGGCTCTCACTGCTAGATAACCCTTGTAACG |
|  | TUA6pro_XFP_R | CTCGCCCTTGCTCACCATTGTCGTTTCGCGGAAGAAAGAAAG |
| *TUA6* protein coding & terminator | Linker1_TUA6coding_F | GGTGGCAGCGGCGGCAGCATGAGAGAGTGCATTTCGATCCACATTG |
|  | TUA6term_attB2_R | GGGGACCACTTTGTACAAGAAAGCTGGGTTTACTGAAATAATGTTTCAGTGCCATCTTTTGTATC |
| *TUB4* promoter | TUB4pro_attB1_F_2 | GGGGACAAGTTTGTACAAAAAAGCAGGCTCAAACTAGTTTTATTACAAAAATCT |
|  | TUB4pro_XFP_R | CTCGCCCTTGCTCACCATTTTTTTTTTTTTTGGTTTCTTCGTCAAGAGC |
| *TUB4* protein coding & terminator | Linker1_TUB4coding_F | GGTGGCAGCGGCGGCAGCATGAGAGAGATCCTTCATATCCAAGGCG |
|  | TUB4term_attB2_R_2 | GGGGACCACTTTGTACAAGAAAGCTGGGTTCAAAAGTTTAACAAATCCAGAACCGATA |
| sGFP | mNG_F | ATGGTGAGCAAGGGCGAG |
|  | GFP_Linker_R | GCTGCCGCCGCTGCCACCCTTGTACAGCTCGTCCAT |
